# DNA Origami Vesicle Sensors with Triggered Cargo Transfer

**DOI:** 10.1101/2023.11.03.565475

**Authors:** Ece Büber, Renukka Yaadav, Tim Schröder, Henri G. Franquelim, Philip Tinnefeld

**Affiliations:** Department of Chemistry and Center for NanoScience, Ludwig-Maximilians-University, Butenandtstraße 5−13, 81377, Munich, Germany; Interfaculty Centre for Bioactive Matter, Leipzig University, ℅ Deutscher Platz 5 (BBZ), 04109 Leipzig, Germany

**Keywords:** Affinity interactions, DNA origami, lipid vesicles, cargo transfer, single-molecule FRET

## Abstract

Interacting with living systems typically involves the ability to address lipid membranes of cellular systems. The first step of interaction of a nanorobot with a cell will thus be the detection of binding to a lipid membrane. Leveraging the programmable nature of DNA origami, we engineered a biosensor harnessing single-molecule Fluorescence Resonance Energy Transfer (smFRET) as transduction mechanism for precise lipid vesicle detection. The system hinges on a hydrophobic ATTO647N modified single-stranded DNA (ssDNA) leash, protruding from a rectangular DNA origami. In a vesicle-free environment, the ssDNA adopts a coiled stance, ensuring high FRET efficiency. However, upon lipid vesicle binding to cholesterol anchors on the DNA origami, the hydrophobic ATTO647N induces the ssDNA to stretch towards the lipid bilayer, leading to reduced FRET efficiency. The strategic placement of cholesterol anchors further modulates this interaction, affecting the observed FRET populations. Beyond its role as a vesicle sensor, we show targeted cargo transport of the acceptor dye unit to the vesicle. The cargo transport is initiated by vesicle bound DNA and a strand displacement reaction. Our interaction platform opens pathways for innovative interaction such as biosensing and molecular transport with complex biosystems.

## INTRODUCTION

In the rapidly advancing field of nanotechnology, the development of dynamic systems that respond to specific molecular signals is a major goal. These systems, capable of translating molecular behavior into practical applications, have the potential to reshape areas such as biosensing, targeted therapeutics, and precise engineering at the nanoscale. Central to these advancements is the DNA origami technique^1-7^, which offers a reliable and customizable framework for designing nanoscale interactions by having stoichiometric and positional control over the DNA structure and attached functional elements. DNA origami, utilizing the innate programmability of DNA sequences, enables the design and realization of intricate nanostructures with exceptional precision. This unique capability has fostered innovations across nanotechnology, particularly in biosensing.^8-11^ With the ability to design custom sensors tailored for specific molecular targets, DNA origami emerges as a powerful tool to address the challenges posed by complex biological systems.^12-19^

Among the challenges, membrane systems and especially lipid vesicles stand out. These membranous sacs play pivotal roles in diverse cellular functions, from molecular transport and signaling to compartmentalization.^20, 21^ Therefore, detecting and characterizing lipid vesicles is of great importance. Sensors based on DNA origami can offer a subtle understanding of how lipid vesicles behave, with the potential to probe, detect, and even manipulate their activities.^22-27^ Bridging the innovative capabilities of DNA origami to the intricate world of lipid vesicles can provide deeper insights into vesicular behaviors and potentially unlock new therapeutic opportunities.^28-31^

DNA origami has uses beyond just detection. As modern medicine and technology advance, the need for precise and controlled cargo transportation at the nanoscale becomes increasingly evident. From targeted drug delivery to the transfer of specific molecular agents, the ability to move and release cargo with specificity could reshape therapeutic strategies. The concept of molecular constructs for precise cargo transport is now an achievable reality in scientific research. This advancement holds significant implications for healthcare, diagnostics, and materials science. In one of the pioneering works, Douglas et al. developed a DNA nanorobot that delivers payloads to cells and changes its structure to release them, showing promise for targeted cell therapy. ^32^ Thubagere et al. created a self-powered DNA robot with three functional domains that can move across a DNA origami sheet and sort two types of molecular cargoes to their respective destinations using a simple algorithm.^33^ In a more recent work involving lipid vesicles, Baumann et al. created a DNA mesh around lipid vesicles for drug delivery, releasing the dye calcein upon triggering. This method increased cytotoxicity in HEK293T cells and holds potential for targeted chemotherapy delivery.^34^ In our research, we have developed a multifunctional system that is not only capable of sensing the presence of a vesicle, but also has the potential to deliver cargo directly to the vesicles it is attached to. By leveraging the capabilities of DNA origami, integrating the precision of single-molecule FRET (smFRET), and utilizing specific binding interactions, we have created innovative biosensing and molecular transport systems with dual functionality.

Combining this foundation with our previous experience with DNA origami nanosensors for lipid vesicle characterization^27^, herein we introduce a DNA origami biosensor tailored for lipid vesicle detection. This system utilizes an ATTO647N labeled single-stranded DNA protrusion, a donor dye ATTO542, and cholesterol anchors. It capitalizes on smFRET and the affinity interactions of ATTO647N, allowing for real-time vesicle sensing and the potential for molecular cargo transport. In our system, ATTO647N serves a dual role: not only is it the acceptor dye in the FRET process, but its hydrophobic nature also drives its interaction with lipid bilayers. Our research highlights how this system could change the way we detect and handle these lipid vesicles, setting the stage for innovative uses in biosensing and transporting molecules.

## RESULTS

The vesicle sensor is crafted from a rectangular DNA origami nanostructure^2, 35^ with the dimensions of 70 × 100 nm. It features a 12 nucleotide (nt) long single-stranded DNA (ssDNA) leash, modified with a ATTO647N fluorophore which serves as the main probe for vesicle sensing (Figure 1a). This probe, due to its hydrophobic and cationic properties, anchors itself in phospholipid vesicles, a behavior noted in prior studies.^36-38^ Moreover, the sensor is equipped with an internally labelled ATTO542 fluorophore, serving as a donor for the fluorescence resonance energy transfer (FRET) (Figure 1a and Figure S1). We postulate that the relative position of the acceptor probe will change based on the presence or absence of lipid vesicles (Figure 1b). Specifically, without the vesicles, the ssDNA probe is more likely to coil up, leading to increased FRET. Conversely, with vesicles present, the leash will stretch out and anchor into the vesicles, increasing the gap between the FRET pair and resulting in decreased FRET. For the purpose of anchoring the sensor to modified glass coverslips, biotin attachments are located at the four corners of this DNA origami structure (Figure 1a and Figure S1). Additionally, the sensor comes with four cholesterol-based anchors that are placed parallel to each other in order to facilitate the capture of lipid vesicles (Figure 1a and Figure S1).^39,40^ These can be adjusted to various distances from the primary sensing probe in order to more efficiently capture and sense vesicles of different sizes.

**Figure 1.**
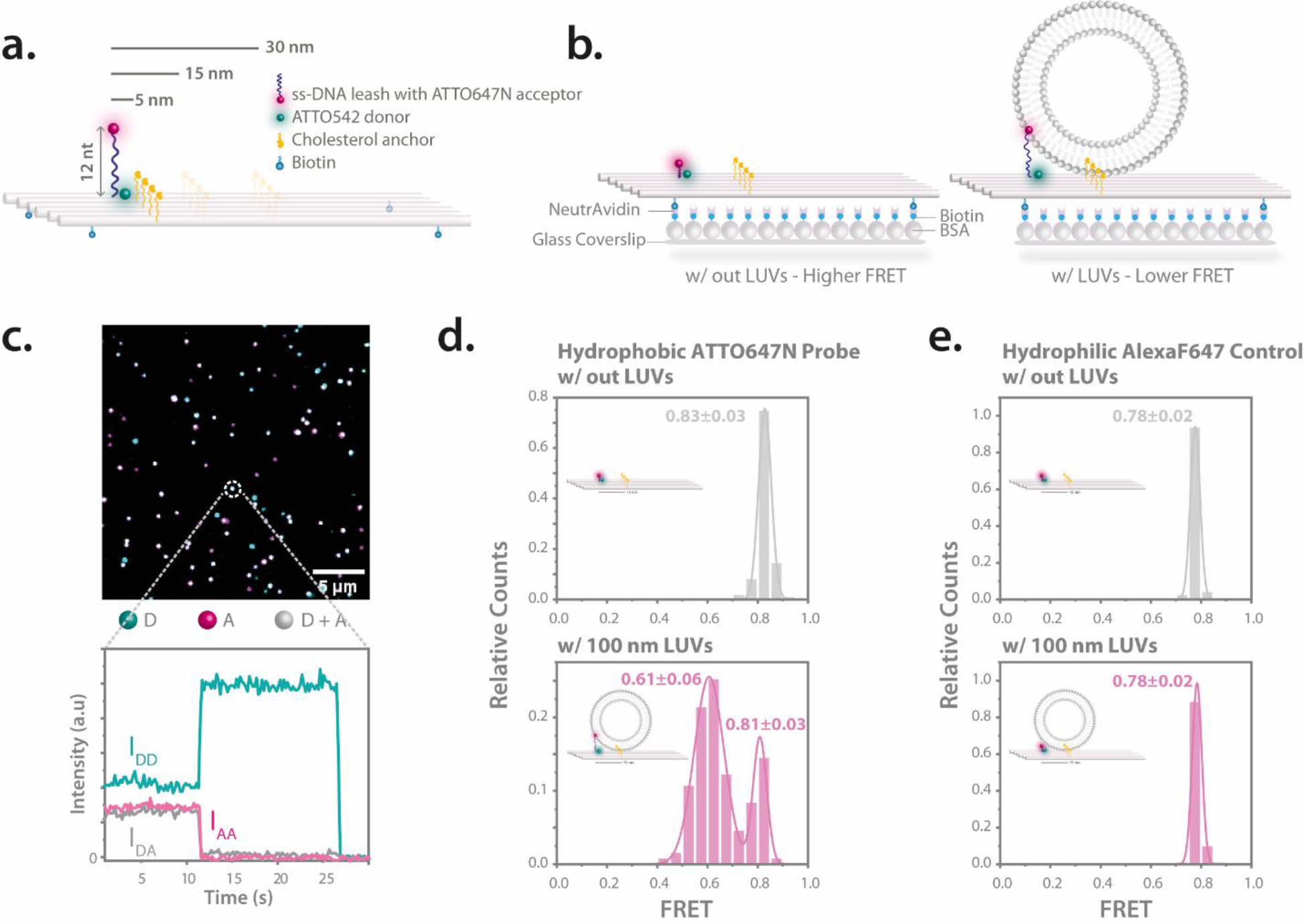
Concept and validation of the vesicle sensor. **(a)** A rectangular DNA origami base equipped with a sensing unit consisting of a ssDNA leash labeled with an ATTO647N acceptor and an ATTO542 donor positioned on the DNA origami base. Cholesterol anchors, strategically placed at various distances for vesicle capture, and biotin moieties for surface attachment are also featured. **(b)** These sensors have the capability to bind to BSA-Biotin-NeutrAvidin-treated glass coverslips using biotin moieties. Without lipid vesicles (left), the ssDNA sensing probe adopts a coiled configuration, positioning it closer to the donor dye, resulting in higher FRET. When lipid vesicles are present (right), the sensing probe elongates to permeate the lipid bilayer, driven by the hydrophobic nature of the ATTO647N fluorophore. Due to the increased distance between the probe and the donor dye, a decreased FRET is observed. **(c)** The image at the top presents a superimposed TIRF image with the donor dye (D) shown in cyan, and the acceptor (A) in magenta.

To visualize individual vesicle sensors, we employed smFRET using total internal reflection fluorescence microscopy (TIRF) on a commercial fluorescence microscope (Nanoimager S, ONI Ltd., UK) with green-red alternating laser excitation (ALEX).^41, 42^ Intensity transients of single spots were extracted from TIRF videos via the iSMS software based on Matlab.^43^ In Figure 1c, a false-color image displays donor dye emission in cyan, acceptor in magenta, and their overlay in gray. We verified the fluorescence as originating from single vesicle sensors by observing single-step photobleaching patterns. For instance, in Figure 1c, the acceptor dye photobleaches around 11 seconds, leading to an increased, unquenched donor fluorescence (I_DD_), while the FRET signal (I_DA_) drops to zero. This synchronized response confirms the presence of a single DNA origami structure exhibiting FRET. We further tracked acceptor emission following its excitation (I_AA_) to study associated photophysical behaviors and prioritize initial acceptor bleaching events. From the intensity data of both the I_DD_ and I_DA_ channels during energy transfer, the FRET efficiency of individual sensors were quantified as

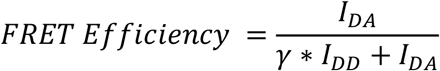

Here, the I_DA_ is corrected to account for the direct excitation of the acceptor at the donor excitation wavelength and for any donor emission leakage into the acceptor emission channel. The γ correction factor compensates for the different quantum yields of the dyes and wavelength-dependent efficiencies of the detection (see SI for detailed materials and methods). For each sample, we then analyzed a minimum of 100 molecules and represented the FRET efficiencies in histograms, complemented with Gaussian fits where relevant.

To validate our design, we contrasted the FRET efficiencies of the sensor both in the absence and presence of lipid vesicles. For this experiment, considering the highest possible overlap with the leash length and 100 nm lipid vesicles, we used the sensors with cholesterol anchors placed at 15 nm distance from the probe position. After immobilizing the sensors on BSA-Biotin-NeutrAvidin-treated glass coverslips, smFRET measurements were performed. In the absence of vesicles, the mean FRET value was 0.83 ± 0.03 (standard deviation of the mean, SD). Upon introducing 1 nM of 100 nm DOPC vesicles with a 1-hour incubation, the mean FRET value shifted to 0.61 ± 0.06 (Figure 1d). This shift confirms the effective binding of lipid vesicle to our sensors and suggests the penetration of hydrophobic ATTO647N into the vesicle membrane. A remaining fraction with a mean FRET of 0.81 ± 0.03 likely represents either sensors devoid of vesicles or those where the probe fails to engage with the vesicles.

To further substantiate the role of affinity interactions in the sensor mechanism, the hydrophobic ATTO647N fluorophore was substituted with AlexaF647 on the ssDNA probe. Previous research has demonstrated that AlexaF647 exhibits minimal interaction with lipid vesicles due to its hydrophilic nature.^36^ When analogous experiments were conducted without and with lipid vesicles for the sensors equipped with AlexaF647, the mean FRET values remained consistent at 0.78 ± 0.02 (Figure 1e). The notably homogenous and narrow distributions for the AlexaF647 probe further underscore the assertion that the distinct behavior of the sensor arises from the hydrophobic affinity interactions between the ATTO647N probe and lipid vesicles.

Gray spots denote sensors that incorporate both the donor and acceptor dyes. An exemplary single-molecule FRET (smFRET) trace at the bottom illustrates the fluorescence intensity over a period, detailing the donor excitation−donor emission (I_DD_) channel (cyan), the donor excitation−acceptor emission (I_DA_) channel (gray), and the acceptor excitation−acceptor emission (I_AA_) channel (magenta). Mean FRET efficiencies are calculated from the I_DD_ and I_DA_ channels. **(d)** FRET efficiency distributions for vesicle sensors in scenarios both without (top) and with (bottom) lipid vesicles. Distributions are shown for the hydrophobic sensing probe with ATTO647N and **(e)** the hydrophilic control probe with AlexaF647. Accompanying illustrations in the plots suggest potential conformations of the probe. The error refers to the standard deviation (SD).

Building on our observations, we investigated the influence of the distance of the FRET probe to the cholesterol anchors. In addition to sensors with cholesterol anchors at 15 nm distance, we assembled vesicle sensors with cholesterol moieties positioned at 5 nm, and 30 nm distance from the probe. Each sensor was subjected to smFRET studies both with and without lipid vesicles.

For the sensor with the cholesterol anchors at a 30 nm distance from the leash, the smFRET results (Figure 2a) reported mean FRET values of 0.81 ± 0.04 and 0.81 ± 0.03 for scenarios without and with vesicles, respectively. The data suggests that, even when vesicles of 100 nm average diameter are present, the probe remains in its coiled state because the vesicles are too distant to be reached by the ATTO647N anchor. This design confirms that the FRET contrast arises solely when the ATTO647N probe anchors into the lipid bilayer—an event only possible when the vesicle is close enough to allow the probe to stretch toward it.

**Figure 2.**
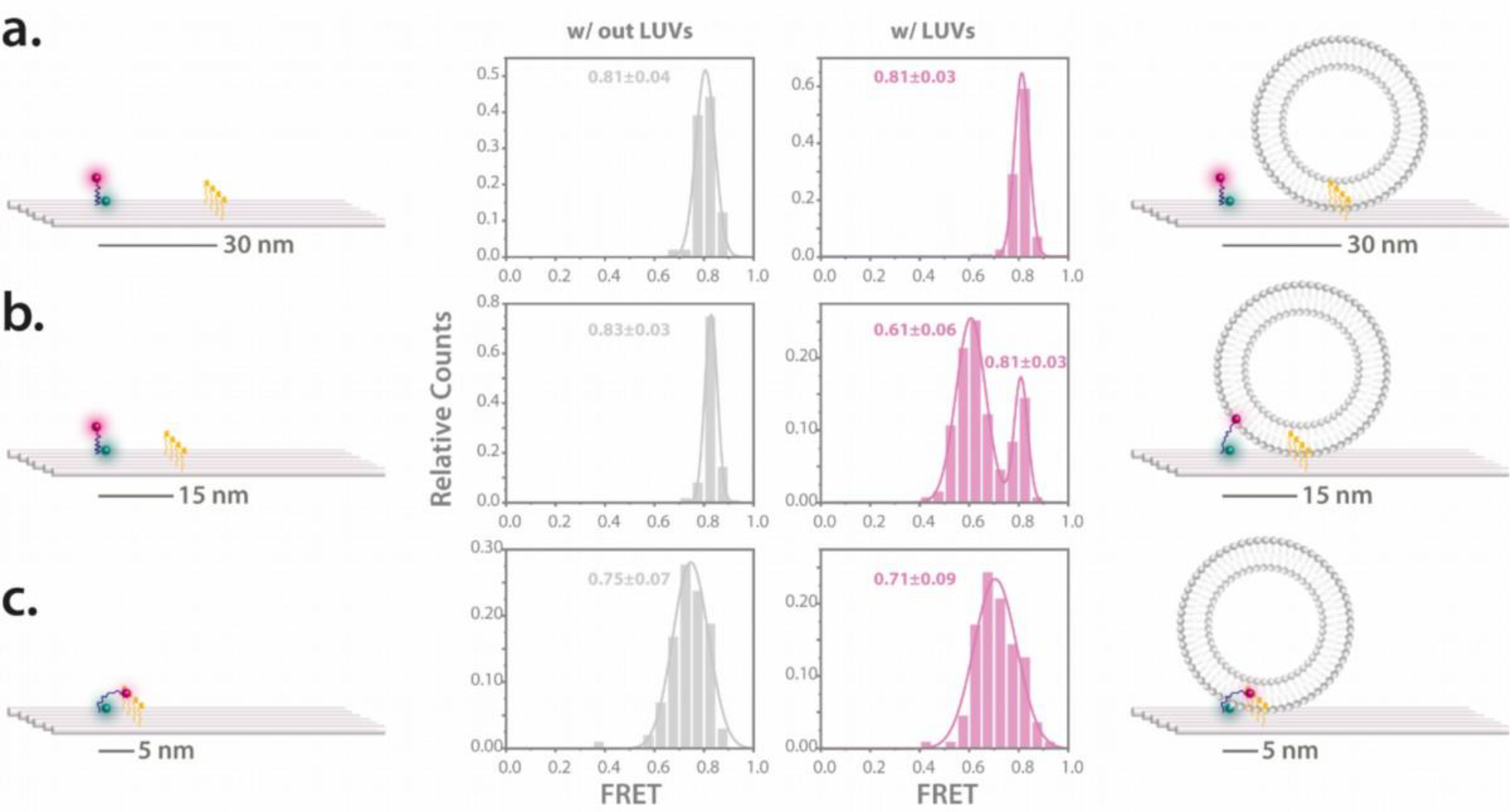
Influence of the cholesterol anchoring distances on the sensor both in the absence and presence of lipid vesicles. The distributions of FRET efficiency for the vesicle sensors, both without and with lipid vesicles, is depicted for **(a)** 30 nm, **(b)** 15 nm, and **(c)** 5 nm cholesterol-FRET probe distances. Illustrations accompanying the data highlight the positions of the cholesterol anchors and potential movement patterns of the sensing probe. The error refers to the standard deviation (SD).

In the sensor variant with 15 nm cholesterols, as already presented in Figure 1d, a clear FRET shift occurred based on vesicle presence, underlining the effect of proximity (Figure 2b). However, the results took a captivating turn when cholesterol anchors were only 5 nm from the probe. Independent of lipid vesicle presence, these sensors exhibited wider FRET distributions with comparable mean values (0.75 ± 0.07 without vesicles and 0.71 ± 0.09 with vesicles, Figure 2c), yet distinct from the other tested distances. This observation can likely be attributed to the strong hydrophobic interactions that allow the ATTO647N probe to already interact with closely positioned cholesterol anchors. Consequently, given the distinctive and reliable FRET shift observed with the 15 nm spacing, we chose to utilize sensors with cholesterol moieties at this distance for the remainder of our study. Additionally, a noteworthy observation was that, upon testing the 5 nm cholesterol configuration with a control system featuring AlexaF647, there was no apparent difference in FRET distributions (Figure S2).

In our pursuit to understand the sensor’s response to varying lipid vesicle sizes, we further examined its behavior with 50 nm, and 200 nm DOPC vesicles using the 15 nm cholesterol as well as a 20 nm cholesterol configuration (Figure S3-a). Notably, while the 15 nm cholesterol-based sensor exhibited minimal variation across vesicle sizes (Figure S3), the 20 nm variant revealed a discernable peak in its interaction with the 200 nm DOPC vesicles, as illustrated in Figure S3-c. This indicates that the positioning of cholesterol in relation to the probe can influence the affinity of the sensor towards differently sized vesicles. Such findings underscore the importance of meticulous design and adaptation in probe creation for vesicle interactions, holding the potential to augment their relevance across diverse scientific and medical arenas.^44-46^ Tailoring probes to align with specific vesicle dimensions can optimize their performance and extend their versatility in multiple disciplines.

Building upon the insights from our prior experiments and recognizing the transformative potential of cargo transport systems in molecular and nanoscale research, we delved deeper into an intricate investigation employing the ATTO647N probe within a strand displacement system. The design approach was systematic. A 17 nt single-stranded DNA was protruding from the probe position on the DNA origami and a 17 nt ATTO647N probe was attached to this protrusion, preserving a 5 nt toehold at the forefront (Figure 3a). The cargo transport system involved a 17 nt cholesterol-modified displacer strand, which has a stronger affinity for the ATTO647N probe. Upon interaction, this displacer strand binds to the probe at the toehold segment, leading to its displacement from the origami structure. More precisely, as we noted higher FRET values without vesicles and a decrease upon vesicle introduction, we further anticipated witnessing a loss of the FRET signal but a persisting colocalization of green and red emission after the displacement of the ATTO647N probe. Due to the reason that a single cholesterol can diffuse in and out of lipid vesicles^47, 48^, the transferred probe can subsequently relocate to surrounding lipid vesicles. Initial smFRET imaging without lipid vesicles revealed a mean FRET value of 0.39 ± 0.04. Presence of lipid vesicles reduced this value to 0.23 ± 0.06, corroborating the consistent working principle of the sensor (Figure 3b and Figure 3c, left and middle panels). The experiment took an intriguing turn when the 17 nt cholesterol-labeled displacer strand was introduced. The FRET signal immediately vanished, yet a red emission persisted (Figure 3b and Figure 3c, right panel). This red emission provided compelling evidence of the probe being successfully detached from the origami and transported to neighbouring lipid vesicles. Another captivating observation was the discrepancy in expected colocalization ratios after the introduction of the displacer strand in the presence of lipid vesicles. Prior to cargo translocation, the system exhibited a high degree of colocalization, as evidenced in Figure 3d, left and middle panels. However, upon introduction of the displacer strand, instead of the anticipated colocalized spots, numerous red-only spots emerged (Figure 3d, right panel). We theorize that this could be attributed to free lipid vesicles adhering to the surface, paired with the mobility of the cholesterol strand across vesicles^47^, causing a transport of the probe from one vesicle to another. This phenomenon may result in diminished colocalization but a surge in red spots. Furthermore, the evident motion in these red spots post-displacement hints at their dynamic diffusion in and across lipid vesicles. Supporting these observations, when lipid vesicles in the system were deliberately ruptured using a 0.05% Tween20 buffer^49^, almost all red signals disappeared, reaffirming the specific vesicle-probe interaction (Figure S4-d). A separate evaluation of the impact of Tween20 on the system showed it only disrupts lipid vesicles and thereby resets the sensor to its original state without vesicle (see Figure S5). To fortify our hypothesis, a control test without vesicles was carried out. Upon introducing the cholesterol-labeled displacer strand, a near-complete disappearance of the red signal was evident which highlights the effectiveness of the strand in displacing the probe (Figure S6). In the absence of lipid vesicles, the displacement process was considerably slower, taking almost an hour for the red signal to fade, whereas in the presence of the vesicles, the red signal faded in just tens of seconds. Supposedly, the cholesterol displacer strand first binds to the lipid vesicles and gets upconcentrated. This upconcentration close to the probe results in increased transfer kinetics. Collectively, these results vividly demonstrate the potential of the vesicle sensor as a robust cargo transport system, emphasizing its versatility and specificity in molecular interactions.

**Figure 3.**
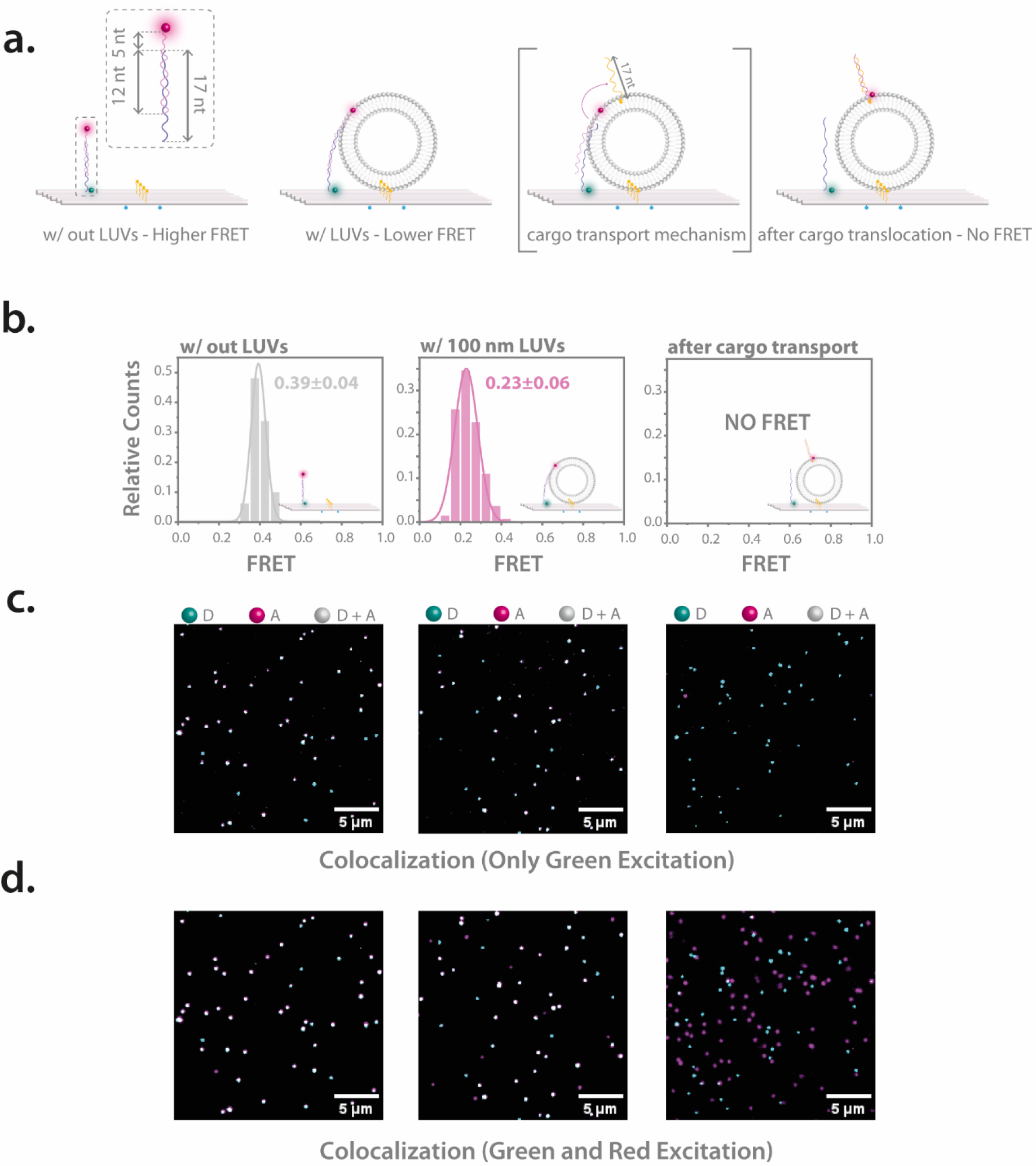
Investigation of cargo translocation to the lipid vesicles. **(a)** Schematic illustrations detailing the cargo transport system. A 17 nt strand protrudes from the DNA origami base, binding to a complementary 17 nt strand labeled with ATTO647N, leaving a 5 nt toehold exposed. The system initially exhibits higher FRET levels in the absence of lipid vesicles. Upon interaction with lipid vesicles, the ATTO647N-labeled strand anchors into the lipid bilayer, leading to reduced FRET. After cargo translocation by strand displacement, the system shows no FRET due to increased separation between the FRET pair. **(b)** FRET histograms displaying the FRET characteristics of the system under different conditions: without lipid vesicles (left), with lipid vesicles (middle), and post-cargo translocation (right), where no FRET is observed. **(c)** Superimposed TIRF images after green-only excitation. The system without vesicles (left) and with vesicles (middle) show clear FRET, evidenced by colocalization. After cargo translocation (right), only green spots are observed, indicating the loss of FRET and absence of colocalization. **(d)** Superimposed TIRF images after both green and red excitation. In the absence (left) and presence (middle) of lipid vesicles, high degree of colocalization is observed. Following cargo translocation (right), the colocalization is lost, coinciding with the absence of FRET. However, numerous red spots remain distributed wherever lipid vesicles are present.

## CONCLUSION

Understanding the interactions between vesicles and probes is not just of biophysical interest— it is a pathway to improving applications in medicine, diagnostics, and molecular biology. Built on the principles of affinity interactions, in this research, we have successfully developed a biosensor predicated on the flexibility and programmability of DNA origami structures. The vesicle sensor utilizes single-molecule Fluorescence Resonance Energy Transfer (smFRET) for the precise detection of lipid vesicles. Central to the effectiveness of the sensor is the hydrophobic ATTO647N dye-modified DNA leash, whose conformational shifts in the presence or absence of lipid vesicles facilitate distinguishable FRET signals. This observed versatility in response is further augmented by the strategic positioning of cholesterol anchors, underscoring the pivotal role of precise molecular design in dictating vesicle-sensor interactions. Moreover, the dynamic behavior of the sensor, as evidenced by the FRET shift upon vesicle interaction and the subsequent engagement of the hydrophobic probe with the vesicle membrane, showcases its potential as a robust mechanism for targeted cargo transport at the nanoscale. The ability to modulate this transport and sensor response based on the proximity of cholesterol anchors provides insights into tailoring DNA origami systems for specific applications, offering a promising avenue for enhanced molecular specificity and efficiency. Our findings on the different affinities of hydrophobic and hydrophilic dyes to lipid vesicles further amplify the importance of selecting appropriate probes for specific biosensing endeavors. Such insights inform future design considerations, optimizing probe interactions for targeted applications. Furthermore, the study of the vesicle sensor within a strand displacement system illuminated the potential of DNA origami structures in molecular transport. Our design adds a physical dimension to conditional cargo transport that first requires the sensing of the vesicle depending on whether it can bridge from binding to the sensing unit followed by cargo transfer through strand displacement which is a different dimension compared to delivery upon molecular logics such as a logic and gate in the presence of two chemical receptors.^32^

In essence, our work has carved a significant stride in the domain of DNA origami-based biosensing. The interplay of FRET signals, hydrophobic dyes, and vesicle presence paints a picture of a sensor system that is not only versatile but is also specific, providing paths for real-world applications. As we advance into an era increasingly reliant on nanotechnology and molecular precision, tools such as the vesicle sensor developed in this study will play an indispensable role. Their versatility, as highlighted in both biosensing and molecular transport, provides a testament to the transformative potential of DNA origami in shaping the future of nanoscale research.

## ASSOCIATED CONTENT

The Supporting Information is available. List of buffers, list of DNA oligonucleotides, materials and methods, and the details of DNA origami nanostructures, sample preparation, characterization techniques, imaging and data analysis.

## ACKNOWLEDGMENT

The authors thank Thomas Bein (Ludwig-Maximilians-University, Department of Chemistry, Munich, Germany) for providing access to their facilities, especially to the dynamic light scattering. E.B. thanks Lorena Manzanares, and Viktorija Glembockytė for the fruitful discussions. This work was funded by the Deutsche Forschungsgemeinschaft (DFG, German Research Foundation) – Project-ID 201269156 – SFB 1032 (A13) and the Bavarian Ministry of Science and the Arts through the ONE MUNICH Project “Munich Multiscale Biofabrication”.

## REFERENCES

1. Seeman, N. C.; Belcher, A. M., Emulating biology: Building nanostructures from the bottom up. Proceedings of the National Academy of Sciences 2002, 99 (suppl_2), 6451–6455.

2. Rothemund, P. W. K., Folding DNA to create nanoscale shapes and patterns. Nature 2006, 440 (7082), 297–302.

3. Douglas, S. M.; Dietz, H.; Liedl, T.; Högberg, B.; Graf, F.; Shih, W. M., Self-assembly of DNA into nanoscale three-dimensional shapes. Nature 2009, 459 (7245), 414–418.

4. Douglas, S. M.; Marblestone, A. H.; Teerapittayanon, S.; Vazquez, A.; Church, G. M.; Shih, W. M., Rapid prototyping of 3D DNA-origami shapes with caDNAno. Nucleic Acids Research 2009, 37 (15), 5001–5006.

5. Castro, C. E.; Kilchherr, F.; Kim, D.-N.; Shiao, E. L.; Wauer, T.; Wortmann, P.; Bathe, M.; Dietz, H., A primer to scaffolded DNA origami. Nature Methods 2011, 8 (3), 221–229.

6. Wagenbauer, K. F.; Engelhardt, F. A. S.; Stahl, E.; Hechtl, V. K.; Stömmer, P.; Seebacher, F.; Meregalli, L.; Ketterer, P.; Gerling, T.; Dietz, H., How We Make DNA Origami. ChemBioChem 2017, 18 (19), 1873–1885.

7. Dey, S.; Fan, C.; Gothelf, K. V.; Li, J.; Lin, C.; Liu, L.; Liu, N.; Nijenhuis, M. A. D.; Saccà, B.; Simmel, F. C.; Yan, H.; Zhan, P., DNA origami. Nature Reviews Methods Primers 2021, 1 (1).

8. Surana, S.; Shenoy, A. R.; Krishnan, Y., Designing DNA nanodevices for compatibility with the immune system of higher organisms. Nature Nanotechnology 2015, 10 (9), 741–747.

9. Xiao, M.; Lai, W.; Man, T.; Chang, B.; Li, L.; Chandrasekaran, A. R.; Pei, H., Rationally Engineered Nucleic Acid Architectures for Biosensing Applications. Chemical Reviews 2019, 119 (22), 11631–11717.

10. Wang, S.; Zhou, Z.; Ma, N.; Yang, S.; Li, K.; Teng, C.; Ke, Y.; Tian, Y., DNA Origami-Enabled Biosensors. Sensors 2020, 20 (23), 6899.

11. Shen, L.; Wang, P.; Ke, Y., DNA Nanotechnology-Based Biosensors and Therapeutics. Advanced Healthcare Materials 2021, 2002205.

12. Koirala, D.; Shrestha, P.; Emura, T.; Hidaka, K.; Mandal, S.; Endo, M.; Sugiyama, H.; Mao, H., Single-Molecule Mechanochemical Sensing Using DNA Origami Nanostructures. Angewandte Chemie International Edition 2014, 53 (31), 8137–8141.

13. Zhang, H.; Chao, J.; Pan, D.; Liu, H.; Qiang, Y.; Liu, K.; Cui, C.; Chen, J.; Huang, Q.; Hu, J.; Wang, L.; Huang, W.; Shi, Y.; Fan, C., DNA origami-based shape IDs for single-molecule nanomechanical genotyping. Nature Communications 2017, 8 (1), 14738.

14. Dutta, P. K.; Zhang, Y.; Blanchard, A. T.; Ge, C.; Rushdi, M.; Weiss, K.; Zhu, C.; Ke, Y.; Salaita, K., Programmable Multivalent DNA-Origami Tension Probes for Reporting Cellular Traction Forces. Nano Lett 2018, 18 (8), 4803–4811.

15. Kosuri, P.; Altheimer, B. D.; Dai, M.; Yin, P.; Zhuang, X., Rotation tracking of genome-processing enzymes using DNA origami rotors. Nature 2019, 572 (7767), 136–140.

16. Raveendran, M.; Lee, A. J.; Sharma, R.; Wälti, C.; Actis, P., Rational design of DNA nanostructures for single molecule biosensing. Nature Communications 2020, 11 (1).

17. Pfeiffer, M.; Trofymchuk, K.; Ranallo, S.; Ricci, F.; Steiner, F.; Cole, F.; Glembockyte, V.; Tinnefeld, P., Single antibody detection in a DNA origami nanoantenna. iScience 2021, 24 (9), 103072.

18. Liu, S.; Jiang, Q.; Zhao, X.; Zhao, R.; Wang, Y.; Wang, Y.; Liu, J.; Shang, Y.; Zhao, S.; Wu, T.; Zhang, Y.; Nie, G.; Ding, B., A DNA nanodevice-based vaccine for cancer immunotherapy. Nature Materials 2021, 20 (3), 421–430.

19. Williamson, P.; Piskunen, P.; Ijäs, H.; Butterworth, A.; Linko, V.; Corrigan, D. K., Signal Amplification in Electrochemical DNA Biosensors Using Target-Capturing DNA Origami Tiles. ACS Sensors 2023, 8 (4), 1471–1480.

20. Dimova, R., Giant Vesicles and Their Use in Assays for Assessing Membrane Phase State, Curvature, Mechanics, and Electrical Properties. Annual Review of Biophysics 2019, 48 (1), 93–119.

21. Tenchov, R.; Bird, R.; Curtze, A. E.; Zhou, Q., Lipid Nanoparticles─From Liposomes to mRNA Vaccine Delivery, a Landscape of Research Diversity and Advancement. ACS Nano 2021, 15 (11), 16982–17015.

22. Czogalla, A.; Kauert, D. J.; Franquelim, H. G.; Uzunova, V.; Zhang, Y.; Seidel, R.; Schwille, P., Amphipathic DNA Origami Nanoparticles to Scaffold and Deform Lipid Membrane Vesicles. Angewandte Chemie International Edition 2015, 54 (22), 6501–6505.

23. Franquelim, H. G.; Khmelinskaia, A.; Sobczak, J.-P.; Dietz, H.; Schwille, P., Membrane sculpting by curved DNA origami scaffolds. Nature Communications 2018, 9 (1).

24. Journot, C. M. A.; Ramakrishna, V.; Wallace, M. I.; Turberfield, A. J., Modifying Membrane Morphology and Interactions with DNA Origami Clathrin-Mimic Networks. ACS Nano 2019, 13 (9), 9973–9979.

25. Liu, L.; Xiong, Q.; Xie, C.; Pincet, F.; Lin, C., Actuating tension-loaded DNA clamps drives membrane tubulation. Science Advances 2022, 8 (41), eadd1830.

26. Hao, P.; Niu, L.; Luo, Y.; Wu, N.; Zhao, Y., Surface Engineering of Lipid Vesicles Based on DNA Nanotechnology. ChemPlusChem 2022, 87 (5).

27. Büber, E.; Schröder, T.; Scheckenbach, M.; Dass, M.; Franquelim, H. G.; Tinnefeld, P., DNA Origami Curvature Sensors for Nanoparticle and Vesicle Size Determination with Single-Molecule FRET Readout. ACS Nano 2023, 17 (3), 3088–3097.

28. Suzuki, Y.; Endo, M.; Sugiyama, H., Mimicking Membrane-Related Biological Events by DNA Origami Nanotechnology. ACS Nano 2015, 9 (4), 3418–3420.

29. Shen, Q.; Grome, M. W.; Yang, Y.; Lin, C., Engineering Lipid Membranes with Programmable DNA Nanostructures. Advanced Biosystems 2020, 4 (1), 1900215.

30. Rubio-Sánchez, R.; Fabrini, G.; Cicuta, P.; Di Michele, L., Amphiphilic DNA nanostructures for bottom-up synthetic biology. Chemical Communications 2021.

31. Kong, Y.; Du, Q.; Li, J.; Xing, H., Engineering bacterial surface interactions using DNA as a programmable material. Chemical Communications 2022, 58 (19), 3086–3100.

32. Douglas, S. M.; Bachelet, I.; Church, G. M., A Logic-Gated Nanorobot for Targeted Transport of Molecular Payloads. Science 2012, 335 (6070), 831–834.

33. Thubagere, A. J.; Li, W.; Johnson, R. F.; Chen, Z.; Doroudi, S.; Lee, Y. L.; Izatt, G.; Wittman, S.; Srinivas, N.; Woods, D.; Winfree, E.; Qian, L., A cargo-sorting DNA robot. Science 2017, 357 (6356), eaan6558.

34. Baumann, K. N.; Schröder, T.; Ciryam, P. S.; Morzy, D.; Tinnefeld, P.; Knowles, T. P. J.; Hernández-Ainsa, S., DNA–Liposome Hybrid Carriers for Triggered Cargo Release. ACS Applied Bio Materials 2022, 5 (8), 3713–3721.

35. Schmied, J. J.; Gietl, A.; Holzmeister, P.; Forthmann, C.; Steinhauer, C.; Dammeyer, T.; Tinnefeld, P., Fluorescence and super-resolution standards based on DNA origami. Nature Methods 2012, 9 (12), 1133–1134.

36. Zhang, Z.; Yomo, D.; Gradinaru, C., Choosing the right fluorophore for single-molecule fluorescence studies in a lipid environment. Biochimica et Biophysica Acta (BBA) - Biomembranes 2017, 1859 (7), 1242–1253.

37. Mobarak, E.; Javanainen, M.; Kulig, W.; Honigmann, A.; Sezgin, E.; Aho, N.; Eggeling, C.; Rog, T.; Vattulainen, I., How to minimize dye-induced perturbations while studying biomembrane structure and dynamics: PEG linkers as a rational alternative. Biochim Biophys Acta Biomembr 2018, 1860 (11), 2436–2445.

38. Ochmann, S. E.; Joshi, H.; Büber, E.; Franquelim, H. G.; Stegemann, P.; Saccà, B.; Keyser, U. F.; Aksimentiev, A.; Tinnefeld, P., DNA Origami Voltage Sensors for Transmembrane Potentials with Single-Molecule Sensitivity. Nano Letters 2021.

39. Langecker, M.; Arnaut, V.; Martin, T. G.; List, J.; Renner, S.; Mayer, M.; Dietz, H.; Simmel, F. C., Synthetic Lipid Membrane Channels Formed by Designed DNA Nanostructures. Science 2012, 338 (6109), 932–936.

40. List, J.; Weber, M.; Simmel, F. C., Hydrophobic Actuation of a DNA Origami Bilayer Structure. Angewandte Chemie International Edition 2014, 53 (16), 4236–4239.

41. Kapanidis, A. N.; Laurence, T. A.; Lee, N. K.; Margeat, E.; Kong, X.; Weiss, S., Alternating-Laser Excitation of Single Molecules. Accounts of Chemical Research 2005, 38 (7), 523–533.

42. Lee, N. K.; Kapanidis, A. N.; Wang, Y.; Michalet, X.; Mukhopadhyay, J.; Ebright, R. H.; Weiss, S., Accurate FRET Measurements within Single Diffusing Biomolecules Using Alternating-Laser Excitation. Biophysical Journal 2005, 88 (4), 2939–2953.

43. Preus, S.; Noer, S. L.; Hildebrandt, L. L.; Gudnason, D.; Birkedal, V., iSMS: single-molecule FRET microscopy software. Nature Methods 2015, 12 (7), 593–594.

44. Khmelinskaia, A.; Mücksch, J.; Petrov, E. P.; Franquelim, H. G.; Schwille, P., Control of Membrane Binding and Diffusion of Cholesteryl-Modified DNA Origami Nanostructures by DNA Spacers. Langmuir 2018, 34 (49), 14921–14931.

45. Ohmann, A.; Göpfrich, K.; Joshi, H.; Thompson, R. F.; Sobota, D.; Ranson, N. A.; Aksimentiev, A.; Keyser, U. F., Controlling aggregation of cholesterol-modified DNA nanostructures. Nucleic Acids Research 2019, 47 (21), 11441–11451.

46. Singh, J. K. D.; Darley, E.; Ridone, P.; Gaston, J. P.; Abbas, A.; Wickham, S. F. J.; Baker, M. A. B., Binding of DNA origami to lipids: maximizing yield and switching via strand displacement. Nucleic Acids Res 2021, 49 (19), 10835–10850.

47. Pfeiffer, I.; Höök, F., Bivalent Cholesterol-Based Coupling of Oligonucletides to Lipid Membrane Assemblies. Journal of the American Chemical Society 2004, 126 (33), 10224–10225.

48. Stengel, G.; Simonsson, L.; Campbell, R. A.; Höök, F., Determinants for Membrane Fusion Induced by Cholesterol-Modified DNA Zippers. The Journal of Physical Chemistry B 2008, 112 (28), 8264–8274.

49. Dresser, L.; Graham, S. P.; Miller, L. M.; Schaefer, C.; Conteduca, D.; Johnson, S.; Leake, M. C.; Quinn, S. D., Tween-20 Induces the Structural Remodeling of Single Lipid Vesicles. The Journal of Physical Chemistry Letters 2022, 13 (23), 5341–5350.

