## Supplementary material for "DNA Origami Vesicle Sensors with Triggered Cargo Transfer": Supplemantary Information

### 1. List of Buffers

| Buffer | Recipe |
|---|---|
| FOB12.5 | 10 mM Tris-HCl, 1 mM EDTA, 12.5 mM $MgCl_2$ |
| PEG Buffer | 12% PEG-8000 (w/v), 10 mM Tris, 1 mM EDTA, 500 mM NaCl, 12 mM $MgCl_2$, pH 7.5 |
| LUV Buffer | 5 mM Tris-HCl, 1 mM EDTA, 0.5 mM Trolox and 650 mM NaCl, pH 7.0 |
| AlexaF-Buffer | 10 mM Tris-HCl, 1 mM EDTA,1 % (wt/v) D-(+)-glucose, 165 units/mL glucose oxidase, 2170 units/mL catalase, 1 mM Trolox, 12.5 mM $MgCl_2$ |

### 2. List of DNA Oligonucleotides

| **5' position** | **Oligonucleotide Sequence** | **Comments** |
|---|---|---|
| | **Sensing Unit Staples** | |
| 12[79] | AAATTAAGTTGACCATTAGATACTTTTGCG**AAAAAAAAAAAA-ATTO647N** | Sensing probe – ATTO647N |
| 12[79] | AAATTAAGTTGACCATTAGATACTTTTGCG**AAAAAAAAAAAA-Alexa647** | Control probe – AlexaF647 |
| 10[79] | **ATTO542-**GATGGCTTATCAAAAAGATTAAGAGCGTCC | Donor dye – ATTO542 |
| | **Anchoring Staples – for Cholesterol labeling** | |
| 11[96] | **TCCTCTACCACCTACATCAC**AATGGTCAACAGGCAAGGCAAAGAG TAATGTG | 5nm chol-1 |
| 13[96] | **TCCTCTACCACCTACATCAC**TAGGTAAACTATTTTTGAGAGATCAA ACGTTA | 5nm chol-2 |
| 9[96] | **TCCTCTACCACCTACATCAC**CGAAAGACTTTGATAAGAGGTCATA TTTCGCA | 5nm chol-3 |
| 7[96] | **TCCTCTACCACCTACATCAC**TAAGAGCAAATGTTTAGACTGGATA GGAAGCC | 5nm chol-4 |
| 10[111] | **TCCTCTACCACCTACATCAC**TTGCTCCTTTCAAATATCGCGTTTGA GGGGGT | 10nm chol-1 |
| 12[111] | **TCCTCTACCACCTACATCAC**TAAATCATATAACCTGTTTAGCTAAC CTTTAA | 10nm chol-2 |
| 14[111] | **TCCTCTACCACCTACATCAC**GAGGGTAGGATTCAAAAGGGTGAGA CATCCAA | 10nm chol-3 |
| 8[111] | **TCCTCTACCACCTACATCAC**AATAGTAAACACTATCATAACCCTC ATTGTGA | 10nm chol-4 |

| | | |
|---|---|---|
| 13[128] | TCCTCTACCACCTACATCACGAGACAGCTAGCTGATAAATTAATTTTTGT | 15nm chol-1 |
| 7[128] | TCCTCTACCACCTACATCACAGACGACAAAGAAGTTTTGCCATAATTCGA | 15nm chol-2 |
| 11[128] | TCCTCTACCACCTACATCACTTTGGGGATAGTAGTAGCATTAAAAGGCCG | 15nm chol-3 |
| 9[128] | TCCTCTACCACCTACATCACGCTTCAATCAGGATTAGAGAGTTATTTTCA | 15nm chol-4 |
| 10[143] | TCCTCTACCACCTACATCACCCAACAGGAGCGAACCAGACCGGAGCCTTTAC | 20nm chol-1 |
| 12[143] | TCCTCTACCACCTACATCACTTCTACTACGCGAGCTGAAAAGGTTACCGCGC | 20nm chol-2 |
| 14[143] | TCCTCTACCACCTACATCACCAACCGTTTCAAATCACCATCAATTCGAGCCA | 20nm chol-3 |
| 8[143] | TCCTCTACCACCTACATCACCTTTTGCAGATAAAAACCAAAATAAAGACTCC | 20nm chol-4 |
| | Chol-GTGATGTAGGTGGTAGAGGA | Cholesterol label on 5’ end |
| | **Surface Binding Staples – for Biotin labeling** | |
| 4[63] | ATAAGGGAACCGGATATTCATTACGTCAGGACGTTGGGAA**GAGGCAATGGCTTGACTCGA** | Biotin-1 |
| 4[255] | AGCCACCACTGTAGCGCGTTTTCAAGGGAGGGAAGGTAAA**GAGGCAATGGCTTGACTCGA** | Biotin-2 |
| 16[63] | CGGATTCTGACGACAGTATCGGCCGCAAGGCGATTAAGTT**GAGGCAATGGCTTGACTCGA** | Biotin-3 |
| 16[255] | GAGAAGAGATAACCTTGCTTCTGTTCGGGAGAAACAATAA**GAGGCAATGGCTTGACTCGA** | Biotin-4 |
| | TCGAGTCAAGCCATTGCCTC-**Bio** | Biotin label on 3’ end |
| | **Strand Displacement System Staples** | |
| 12[79] | AAATTAAGTTGACCATTAGATACTTTTGCG**GGGAAGTTCCAGCAGGG** | 17nt protrusion at the probe position |
| | GTTCCAGCAGGGATTCA-**Chol** | Cholesterol label on 3’ end |
| | **ATTO647N-**TGAATCCCTGCTGGAAC | Sensing probe for the strand displacement system, 5’ ATTO647N label |
| | **Unmodified Staples** | |
| 20[207] | GCGGAACATCTGAATAATGGAAGGTACAAAAT | |
| 23[192] | ACCCTTCTGACCTGAAAGCGTAAGACGCTGAG | |
| 0[175] | TCCACAGACAGCCCTCATAGTTAGCGTAACGA | |
| 1[256] | CAGGAGGTGGGGTCAGTGCCTTGAGTCTCTGAATTTACCG | |
| 21[64] | GCCCTTCAGAGTCCACTATTAAAGGGTGCCGT | |
| 16[207] | ACCTTTTTATTTTAGTTAATTTCATAGGGCTT | |

| | |
|---|---|
| 21[128 ] | GCGAAAAATCCCTTATAAATCAAGCCGGCG |
| 18[271 ] | CTTTTACAAAATCGTCGCTATTAGCGATAG |
| 8[239] | AAGTAAGCAGACACCACGGAATAATATTGACG |
| 1[192] | GCGGATAACCTATTATTCTGAAACAGACGATT |
| 2[239] | GCCCGTATCCGGAATAGGTGTATCAGCCCAAT |
| 14[271 ] | TTAGTATCACAATAGATAAGTCCACGAGCA |
| 0[239] | AGGAACCCATGTACCGTAACACTTGATATAA |
| 9[224] | AAAGTCACAAAATAAACAGCCAGCGTTTTA |
| 20[175 ] | ATTATCATTCAATATAATCCTGACAATTAC |
| 6[175] | CAGCAAAAGGAAACGTCACCAATGAGCCGC |
| 12[239 ] | CTTATCATTCCCGACTTGCGGGAGCCTAATTT |
| 3[96] | ACACTCATCCATGTTACTTAGCCGAAAGCTGC |
| 18[79] | GATGTGCTTCAGGAAGATCGCACAATGTGA |
| 6[271] | ACCGATTGTCGGCATTTTCGGTCATAATCA |
| 21[224 ] | CTTTAGGGCCTGCAACAGTGCCAATACGTG |
| 1[96] | AAACAGCTTTTTGCGGGATCGTCAACACTAAA |
| 19[224 ] | CTACCATAGTTTGAGTAACATTTAAAATAT |
| 21[192 ] | TGAAAGGAGCAAATGAAAAATCTAGAGATAGA |
| 18[207 ] | CGCGCAGATTACCTTTTTTAATGGGAGAGACT |
| 4[47] | GACCAACTAATGCCACTACGAAGGGGGTAGCA |
| 13[32] | AACGCAAAATCGATGAACGGTACCGGTTGA |
| 1[160] | TTAGGATTGGCTGAGACTCCTCAATAACCGAT |
| 22[271 ] | CAGAAGATTAGATAATACATTTGTCGACAA |
| 1[32] | AGGCTCCAGAGGCTTTGAGGACACGGGTAA |
| 16[79] | GCGAGTAAAAATATTTAAATTGTTACAAAG |
| 4[239] | GCCTCCCTCAGAATGGAAAGCGCAGTAACAGT |
| 16[143 ] | GCCATCAAGCTCATTTTTTAACCACAAATCCA |
| 4[207] | CCACCCTCTATTCACAAACAAATACCTGCCTA |
| 19[96] | CTGTGTGATTGCGTTGCGCTCACTAGAGTTGC |
| 22[47] | CTCCAACGCAGTGAGACGGGCAACCAGCTGCA |
| 12[47] | TAAATCGGGATTCCCAATTCTGCGATATAATG |
| 8[271] | AATAGCTATCAATAGAAAATTCAACATTCA |
| 7[56] | ATGCAGATACATAACGGGAATCGTCATAAATAAAGCAAAG |
| 18[143 ] | CAACTGTTGCGCCATTCGCCATTCAAACATCA |
| 14[47] | AACAAGAGGGATAAAAATTTTTAGCATAAAGC |
| 14[207 ] | AATTGAGAATTCTGTCCAGACGACTAAACCAA |
| 0[271] | CCACCCTCATTTTCAGGGATAGCAACCGTACT |
| 23[32] | CAAATCAAGTTTTTTGGGGTCGAAACGTGGA |

| | |
|---|---|
| 10[271] | ACGCTAACACCCACAAGAATTGAAAATAGC |
| 3[128] | AGCGCGATGATAAATTGTGTCGTGACGAGA |
| 10[239] | GCCAGTTAGAGGGTAATTGAGCGCTTTAAGAA |
| 5[128] | AACACCAAATTTCAACTTTAATCGTTTACC |
| 0[207] | TCACCAGTACAAACTACAACGCCTAGTACCAG |
| 19[56] | TACCGAGCTCGAATTCGGGAAACCTGTCGTGCAGCTGATT |
| 17[192] | CATTTGAAGGCGAATTATTCATTTTTGTTTGG |
| 4[79] | GCGCAGACAAGAGGCAAAAGAATCCCTCAG |
| 20[239] | ATTTTAAAATCAAAATTATTTGCACGGATTCG |
| 19[160] | GCAATTCACATATTCCTGATTATCAAAGTGTA |
| 15[128] | TAAATCAAAATAATTCGCGTCTCGGAAACC |
| 0[143] | TCTAAAGTTTTGTCGTCTTTCCAGCCGACAA |
| 10[47] | CTGTAGCTTGACTATTATAGTCAGTTCATTGA |
| 17[128] | AGGCAAAGGGAAGGGCGATCGGCAATTCCA |
| 21[96] | AGCAAGCGTAGGGTTGAGTGTTGTAGGGAGCC |
| 2[207] | TTTCGGAAGTGCCGTCGAGAGGGTGAGTTTCG |
| 22[239] | TTAACACCAGCACTAACAACTAATCGTTATTA |
| 19[32] | GTCGACTTCGGCCAACGCGCGGGGTTTTTC |
| 11[64] | GATTTAGTCAATAAAGCCTCAGAGAACCCTCA |
| 6[239] | GAAATTATTGCCTTTAGCGTCAGACCGGAACC |
| 18[47] | CCAGGGTTGCCAGTTTGAGGGGACCCGTGGGA |
| 10[191] | GAAACGATAGAAGGCTTATCCGGTCTCATCGAGAACAAGC |
| 3[160] | TTGACAGGCCACCACCAGAGCCGCGATTTGTA |
| 8[47] | ATCCCCCTATACCACATTCAACTAGAAAAATC |
| 13[64] | TATATTTTGTCATTGCCTGAGAGTGGAAGATTGTATAAGC |
| 5[160] | GCAAGGCCTCACCAGTAGCACCATGGGCTTGA |
| 5[32] | CATCAAGTAAAACGAACTAACGAGTTGAGA |
| 11[224] | GCGAACCTCCAAGAACGGGTATGACAATAA |
| 17[160] | AGAAAACAAAGAAGATGATGAAACAGGCTGCG |
| 4[271] | AAATCACCTTCCAGTAAGCGTCAGTAATAA |
| 15[32] | TAATCAGCGGATTGACCGTAATCGTAACCG |
| 13[184] | GACAAAAGGTAAAGTAATCGCCATATTTAACAAAACTTTT |
| 20[143] | AAGCCTGGTACGAGCCGGAAGCATAGATGATG |
| 14[79] | GCTATCAGAAATGCAATGCCTGAATTAGCA |
| 6[207] | TCACCGACGCACCGTAATCAGTAGCAGAACCG |
| 3[192] | GGCCTTGAAGAGCCACCACCCTCAGAAACCAT |
| 23[256] | CTTTAATGCGCGAACTGATAGCCCCACCAG |
| 2[47] | ACGGCTACAAAAGGAGCCTTTAATGTGAGAAT |

| | |
|---|---|
| 13[224 ] | ACAACATGCCAACGCTCAACAGTCTTCTGA |
| 20[79] | TTCCAGTCGTAATCATGGTCATAAAAGGGG |
| 9[64] | CGGATTGCAGAGCTTAATTGCTGAAACGAGTA |
| 22[175 ] | ACCTTGCTTGGTCAGTTGGCAAAGAGCGGA |
| 23[160 ] | TAAAAGGGACATTCTGGCCAACAAAGCATC |
| 16[239 ] | GAATTTATTTAATGGTTTGAAATATTCTTACC |
| 3[32] | AATACGTTTGAAAGAGGACAGACTGACCTT |
| 23[224 ] | GCACAGACAATATTTTTGAATGGGGTCAGTA |
| 15[192 ] | TCAAATATAACCTCCGGCTTAGGTAACAATTT |
| 7[192] | ATACATACCGAGGAAACGCAATAAGAAGCGCATTAGACGG |
| 12[271 ] | TGTAGAAATCAAGATTAGTTGCTCTTACCA |
| 4[111] | GACCTGCTCTTTGACCCCCAGCGAGGGAGTTA |
| 17[96] | GCTTTCCGATTACGCCAGCTGGCGGCTGTTTC |
| 23[96] | CCCGATTTAGAGCTTGACGGGGAAAAAGAATA |
| 16[111 ] | TGTAGCCATTAAAATTCGCATTAAATGCCGGA |
| 0[79] | ACAACTTTCAACAGTTTCAGCGGATGTATCGG |
| 1[128] | TGACAACTCGCTGAGGCTTGCATTATACCA |
| 6[111] | ATTACCTTTGAATAAGGCTTGCCCAAATCCGC |
| 20[271 ] | CTCGTATTAGAAATTGCGTAGATACAGTAC |
| 22[79] | TGGAACAACCGCCTGGCCCTGAGGCCCGCT |
| 18[239 ] | CCTGATTGCAATATATGTGAGTGATCAATAGT |
| 9[256] | GAGAGATAGAGCGTCTTTCCAGAGGTTTTGAA |
| 22[143 ] | TCGGCAAATCCTGTTTGATGGTGGACCCTCAA |
| 10[207 ] | ATCCCAATGAGAATTAACTGAACAGTTACCAG |
| 21[32] | TTTTCACTCAAAGGGCGAAAAACCATCACC |
| 5[192] | CGATAGCATTGAGCCATTTGGGAACGTAGAAA |
| 8[79] | AATACTGCCCAAAAGGAATTACGTGGCTCA |
| 15[224 ] | CCTAAATCAAAATCATAGGTCTAAACAGTA |
| 0[111] | TAAATGAATTTTCTGTATGGGATTAATTTCTT |
| 6[143] | GATGGTTTGAACGAGTAGTAAATTTACCATTA |
| 13[256 ] | GTTTATCAATATGCGTTATACAAACCGACCGTGTGATAAA |
| 14[239 ] | AGTATAAAGTTCAGCTAATGCAGATGTCTTTC |
| 22[111 ] | GCCCGAGAGTCCACGCTGGTTTGCAGCTAACT |
| 3[224] | TTAAAGCCAGAGCCGCCACCCTCGACAGAA |
| 1[224] | GTATAGCAAACAGTTAATGCCCAATCCTCA |
| 19[128 ] | CACAACAGGTGCCTAATGAGTGCCCAGCAG |

| | |
|---|---|
| 2[79] | CAGCGAAACTTGCTTTCGAGGTGTTGCTAA |
| 2[271] | GTTTTAACTTAGTACCGCCACCCAGAGCCA |
| 7[224] | AACGCAAAGATAGCCGAACAAACCCTGAAC |
| 19[248] | CGTAAAACAGAAATAAAAATCCTTTGCCCGAAAGATTAGA |
| 15[160] | ATCGCAAGTATGTAAATGCTGATGATAGGAAC |
| 6[47] | TACGTTAAAGTAATCTTGACAAGAACCGAACT |
| 18[111] | TCTTCGCTGCACCGCTTCTGGTGCGGCCTTCC |
| 16[271] | CTTAGATTTAAGGCGTTAAATAAAGCCTGT |
| 1[64] | TTTATCAGGACAGCATCGGAACGACACCAACCTAAAACGA |
| 2[143] | ATATTCGGAACCATCGCCCACGCAGAGAAGGA |
| 16[47] | ACAAACGGAAAAGCCCCAAAAACACTGGAGCA |
| 7[32] | TTTAGGACAAATGCTTTAAACAATCAGGTC |
| 15[96] | ATATTTTGGCTTTCATCAACATTATCCAGCCA |
| 20[111] | CACATTAAAATTGTTATCCGCTCATGCGGGCC |
| 2[111] | AAGGCCGCTGATACCGATAGTTGCGACGTTAG |
| 19[192] | ATTATACTAAGAAACCACCAGAAGTCAACAGT |
| 17[224] | CATAAATCTTTGAATACCAAGTGTTAGAAC |
| 21[256] | GCCGTCAAAAAACAGAGGTGAGGCCTATTAGT |
| 0[47] | AGAAAGGAACAACTAAAGGAATTCAAAAAAA |
| 7[248] | GTTTATTTTGTCACAATCTTACCGAAGCCCTTTAATATCA |
| 4[143] | TCATCGCCAACAAAGTACAACGGACGCCAGCA |
| 18[175] | CTGAGCAAAAATTAATTACATTTTGGGTTA |
| 11[256] | GCCTTAAACCAATCAATAATCGGCACGCGCCT |
| 6[79] | TTATACCACCAAATCAACGTAACGAACGAG |
| 22[207] | AGCCAGCAATTGAGGAAGGTTATCATCATTTT |
| 12[207] | GTACCGCAATTCTAAGAACGCGAGTATTATTT |
| 8[207] | AAGGAAACATAAAGGTGGCAACATTATCACCG |
| 5[96] | TCATTCAGATGCGATTTTAAGAACAGGCATAG |
| 21[160] | TCAATATCGAACCTCAAATATCAATTCCGAAA |
| 4[175] | CACCAGAAAGGTTGAGGCAGGTCATGAAAG |
| 23[128] | AACGTGGCGAGAAAGGAAGGGAAACCAGTAA |
| 23[64] | AAAGCACTAAATCGGAACCCTAATCCAGTT |
| 5[224] | TCAAGTTTCATTAAAGGTGAATATAAAAGA |
| 9[32] | TTTACCCCAACATGTTTTAAATTTCCATAT |
| 20[47] | TTAATGAACTAGAGGATCCCCGGGGGGTAACG |
| 11[32] | AACAGTTTTGTACCAAAAACATTTTATTTC |
| 2[175] | TATTAAGAAGCGGGGTTTTGCTCGTAGCAT |

| | |
|---|---|
| 16[175 ] | TATAACTAACAAAGAACGCGAGAACGCCAA |
| 17[32] | TGCATCTTTCCCAGTCACGACGGCCTGCAG |

### 3. Methods

#### 3.1. Design and production of the vesicle sensors

As the base of the vesicle sensors, a rectangular DNA origami based on a 7249 nt long scaffold derived from the M13mp18 bacteriophage was designed in the software CaDNAno[1] with several modifications indicated in Figure S1. All staple strand sequences and modifications are given in the list of DNA oligonucleotides in Section 2.

The design includes positions for the sensing unit involving fluorophore labelled staples, surface binding staples with biotin modifications, and vesicle anchoring staples with cholesterol modifications. High performance liquid chromatograph (HPLC) purified sensing unit oligonucleotides labeled with ATTO647N and ATTO 542 were obtained from Biomers GmbH. High purity salt free (HPSF) purified unmodified staple oligonucleotides and staples for the strand displacement reaction system were supplied from IDT (Integrated DNA Technologies, Inc.). 3' Biotin and 5' Cholesterol-TEG labeled oligonucleotides as well as the control probe carrying AlexaF647 fluorophore were purchased from Eurofins Genomics GmbH.

The DNA origami nanostructures were folded using a one-pot reaction mix in a thermocycler (primus 25, peqlab). Briefly, 25 nM of scaffold DNA, unmodified oligonucleotides at a concentration 10-times, fluorophore-labeled and biotinylated oligonucleotides, both at a concentration 30-times that of the scaffold were mixed in FOB12.5 buffer and subjected to a multistep thermocycling procedure. The folding began by heating the mixture to 70°C, allowing it to equilibrate for 5 minutes, and then gradually cooling it to 20°C at a rate of 1°C per minute.

Once folded, the DNA origami nanostructures were purified using PEG precipitation. To achieve this, the sample was mixed with the PEG buffer in equal parts and then centrifuged for 30 min at 16,000 g in a 4°C environment. The resulting supernatant was removed, leaving the pellet which was further dissolved in FOB12 buffer and an equal amount of PEG buffer was added in the solution. Two times of the PEG purification were performed for each sample. For the purpose of labeling the vesicle sensors with cholesterol anchors, the neatly folded nanostructures were mixed with cholesterol-labeled oligonucleotides (at a 20-fold excess for each labeling position) and left for overnight incubation at room temperature. A subsequent PEG precipitation step was performed for further purification.

#### 3.2. Preparation and characterization of lipid vesicles

All lipids used in the study were obtained from Avanti Polar Lipids, located in Alabaster, AL, USA, unless stated otherwise. In order to prepare large unilamellar vesicles (LUVs) of 100% DOPC (1,2-dioleoyl-sn-glycero-3-phosphocholine), the lipid solutions in chloroform were dried using a nitrogen stream, and the remaining chloroform was evaporated under vacuum for approximately 3 hours in a desiccator. The resulting lipid film was rehydrated in the LUV buffer, resulting in a final lipid concentration of 2.5 mM. To ensure uniform vesicle size, the solution underwent seven freeze-and-thaw cycles alternating between liquid nitrogen and a water bath at 70°C. Subsequently, the vesicles were extruded using a LiposoFast Basic extruder (Avestin, INC.), employing Nucleopore PC membranes (Whatman, Cytiva Ltd.) with the desired pore size.

The size and distribution of the extruded LUV suspensions were evaluated using dynamic light scattering (DLS). These measurements were taken at dilutions that produced attenuation values between 7 and 8. Every sample underwent 10 consecutive measurements after they were allowed to incubate for 60 seconds. These measurements were conducted in disposable polystyrene cuvettes utilizing a Zetasizer Nano ZSP (Malvern Instruments, Worcestershire, UK). The device operated with a 633 nm incident wavelength and detected backscattering at an angle of 173°. Both intensity and number-normalized size distributions for the LUV samples were deduced from the correlation functions. These calculations employed the General Purpose Non-Negative Least Squares (NNLS) algorithm available in the Malvern Zetasizer Software version 7.13.

#### 3.3. Surface preparation

24 x 60 mm microscope coverslips with a thickness of 170 µm utilized in the experiments (Carl Roth GmbH, Germany). Prior to use, they were cleaned with a UV-Ozone cleaner (PSD-UV4,

Novascan Technologies, USA) operating at 100°C for 30 minutes. Once cleaned, they were fitted with self-adhesive 150 µL SecureSeal Hybridization Chambers from Grace Bio-Labs. For the passivation process, the coverslips were treated with biotinylated-BSA (1.0 mg/mL, Thermo Fisher Scientific Inc.), allowing it to incubate for 15 minutes. This was succeeded by an additional functionalization step using NeutrAvidin (1.0 mg/mL, Thermo Fischer Scientific Inc.), with another 15-minute incubation. After each of these steps, the coverslips underwent a triple wash with 1× PBS buffer.

When the glass coverslips are ready for incorporating the sensors, roughly 50 pM of vesicle sensors (diluted in the LUV buffer) were immobilized onto NeutrAvidin-treated glass coverslips using biotin modifications on the sensors.

For tests involving lipid vesicles, vesicles were diluted to a 1 nM concentration using LUV buffer. The sensor-coated coverslips were incubated with the diluted vesicles for around an hour to capture the vesicles via cholesterol modifications. Post this anchoring phase, the coverslips were rinsed thrice with the LUV buffer.

##### 3.4. Imaging and data analysis

The smFRET experiments were executed with a Nanoimager S (ONI Ltd. UK), under Total Internal Reflection Fluorescence (TIRF) illumination. A red excitation wavelength of 638 nm was used, while the green excitation was set at 532 nm. Initially, the objective was adjusted to focus on the sample plane, followed by activating the auto-focus function. The alternating laser excitation setting employed both green and red lasers[2-4], with power outputs of 5 mW and 8 mW, respectively. For each sample, around 10 videos were captured over a field of view spanning 50 µm × 80 µm. Videos consisted of 500 frames, each with an exposure time of 100 ms, leading to a total video duration of 50 seconds.

In all the experiments utilizing the sensing probe with ATTO647N, a standard LUV buffer (5 mM Tris-HCl, 1 mM EDTA, 0.5 mM Trolox and 650 mM NaCl, pH 7.0) was utilized. However, for the experiments involving the control probe with AlexaF647, the buffer composition was specifically altered to enhance the photostability of this fast bleaching fluorophore. The modified AlexaF-buffer incorporated a reducing and oxidizing system (ROXS)[5] along with oxygen scavenging agents. In detail, the imaging buffer for the AlexaF647 experiments consisted of 10 mM Tris-HCl and 1 mM EDTA with 1 % (wt/v) D-(+)-glucose (Sigma Aldrich, USA), 165 units/mL glucose oxidase (G2133, Sigma Aldrich, USA), 2170 units/mL catalase (C3155, Sigma Aldrich, USA), 1 mM Trolox (~12% was converted to Troloxquinone under UV light)[6] and 650 mM NaCl.

Analysis of the FRET data was performed using the iSMS software on MATLAB.[7] The process integrated individual green and red emission channels. The peakfinder algorithm was used to distinguish donor and acceptor peaks, facilitating the detection of FRET pairs. From the captured sequences, intensity-time trajectories of specific immobilized molecules were extracted. These single-molecule data sets were then further examined to detect DNA origami structures manifesting FRET. Interrelations of FRET across different channels, including $I_{DD}$ (donor excitation-donor emission), $I_{DA}$ (donor excitation-acceptor emission), and $I_{AA}$ (acceptor excitation-acceptor emission), were analyzed. Instances where an increase in the $I_{DD}$ channel intensity and a decrease in the $I_{DA}$ channel occurred concurrently with a sudden drop in the $I_{AA}$ channel led to that specific data set being chosen for extended evaluation. Using the intensity data of these channels for each molecule, the average FRET efficiency of every sensor was calculated over the entire energy transfer duration.

$$FRET\ efficiency = \frac{I_{DA}}{\gamma \cdot I_{DD} + I_{DA}}$$

The $I_{DA}$ is corrected for direct acceptor excitation at the donor excitation wavelength and leakage of donor emission into the acceptor emission channel. The corrected $I_{DA}$ is determined as

$$I_{DA} = I_{A,raw} - D_{leakage} * I_{DD} - A_{direct} * I_{AA}$$

where $I_{A,raw}$ represents the total intensity detected in the acceptor emission channel, and $I_{AA}$ signifies the direct excitation of the acceptor.

Wherever applicable, the correction factors γ, $D_{leakage}$ and $A_{direct}$, are calculated as in the following:

The γ correction factor describes the relative difference in the number of photons measured of the acceptor and the donor for the same number of excited states. In iSMS γ is calculated for all molecules in which the acceptor bleaches before the acceptor:

$$\gamma = \frac{I_{DA,1} - I_{DA,2}}{I_{DD,2} - I_{DD,1}}$$

The $D_{leakage}$ accounts for the amount of leakage of the donor emission into the acceptor emission channel upon donor excitation and is calculated as

$$D_{leakage} = avg(\frac{I_{DA}(t)}{I_{DD}(t)})$$

in the time interval after the acceptor has bleached and before the donor has bleached, corresponding to a donor-only signal.

The $A_{direct}$ accounts for the direct excitation of the acceptor at the donor wavelength and is calculated as

$$A_{direct} = avg(\frac{I_{DA}(t)}{I_{AA}(t)})$$

in the time interval after the donor has bleached and before the acceptor has bleached. From individual transients, we calculated average values for all the correction factors and used them for FRET efficiency calculations.

### 4. Figures

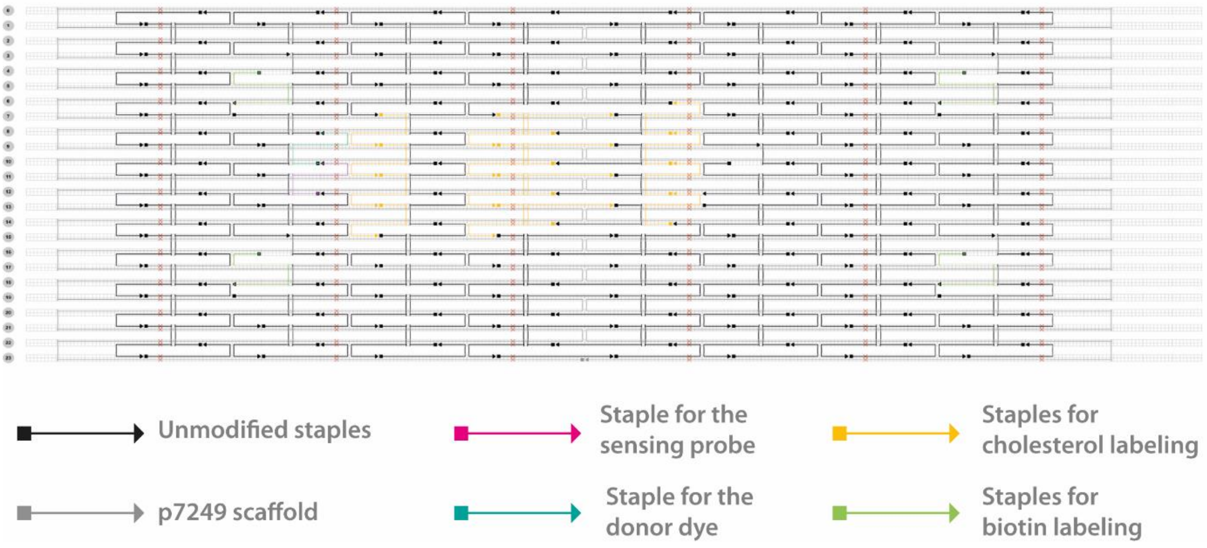

**Figure S1.** CaDNAno design of the vesicle sensor. Zoom in to see details. Sensing probe position is colored in magenta, and the donor dye position is in cyan. Positions of the staples for cholesterol modification are colored in orange, and for biotin modification are in green.

Unmodified staples are colored in black, and M13 p7249 scaffold is colored in gray. The sequences and details of all the staples incorporated in the DNA origami design are listed in Section 2.

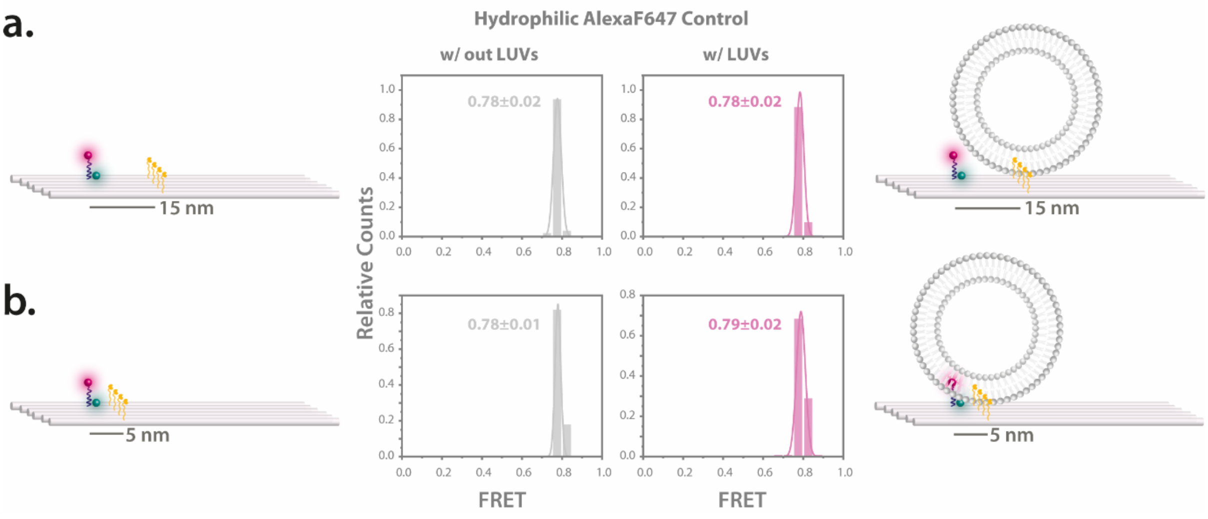

**Figure S2.** Sensor with the control probe AlexaF647. Effect of different cholesterol anchoring distances both in the absence and presence of lipid vesicles. The distribution of FRET efficiency for the vesicle sensors, both without and with lipid vesicles, is depicted for **(a)** 15 nm, and **(b)** 5 nm cholesterol configurations. Illustrations accompanying the data highlight the positions of the cholesterol anchors and potential movement patterns of the sensing probe. The error refers to the standard deviation (SD).

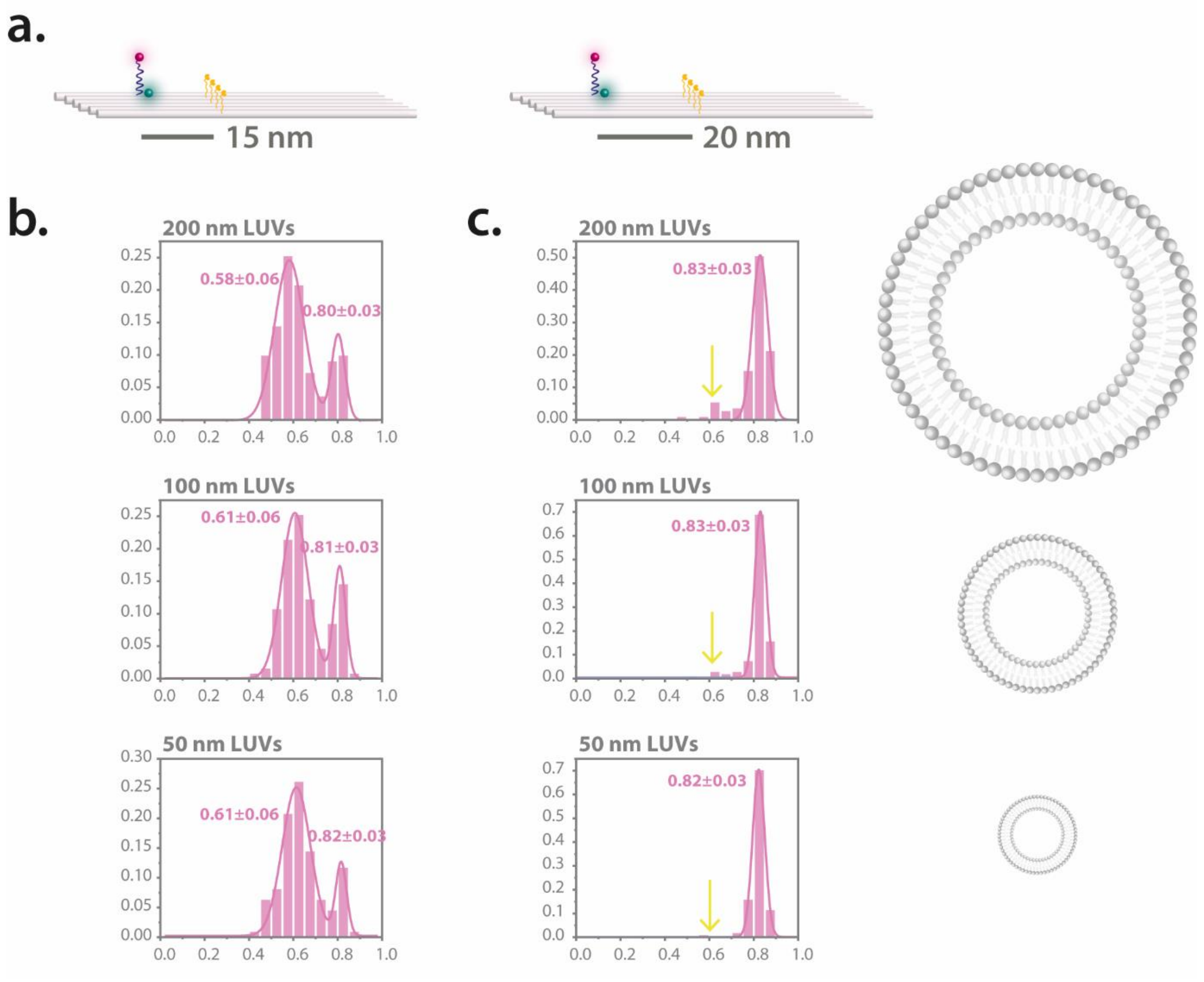

**Figure S3.** Sensor with different sizes of vesicles. **(a)** The sketches of the utilized sensors showing the position of the anchors. FRET efficiency distributions of the vesicle sensors **(b)** with 15 nm and **(c)** with 20 nm cholesterol configurations tested with lipid vesicles of 200 nm, 100 nm and 50 nm (from top to bottom). The yellow arrows highlight the slight increase in the lower FRET population for larger vesicles. The error refers to the standard deviation (SD).

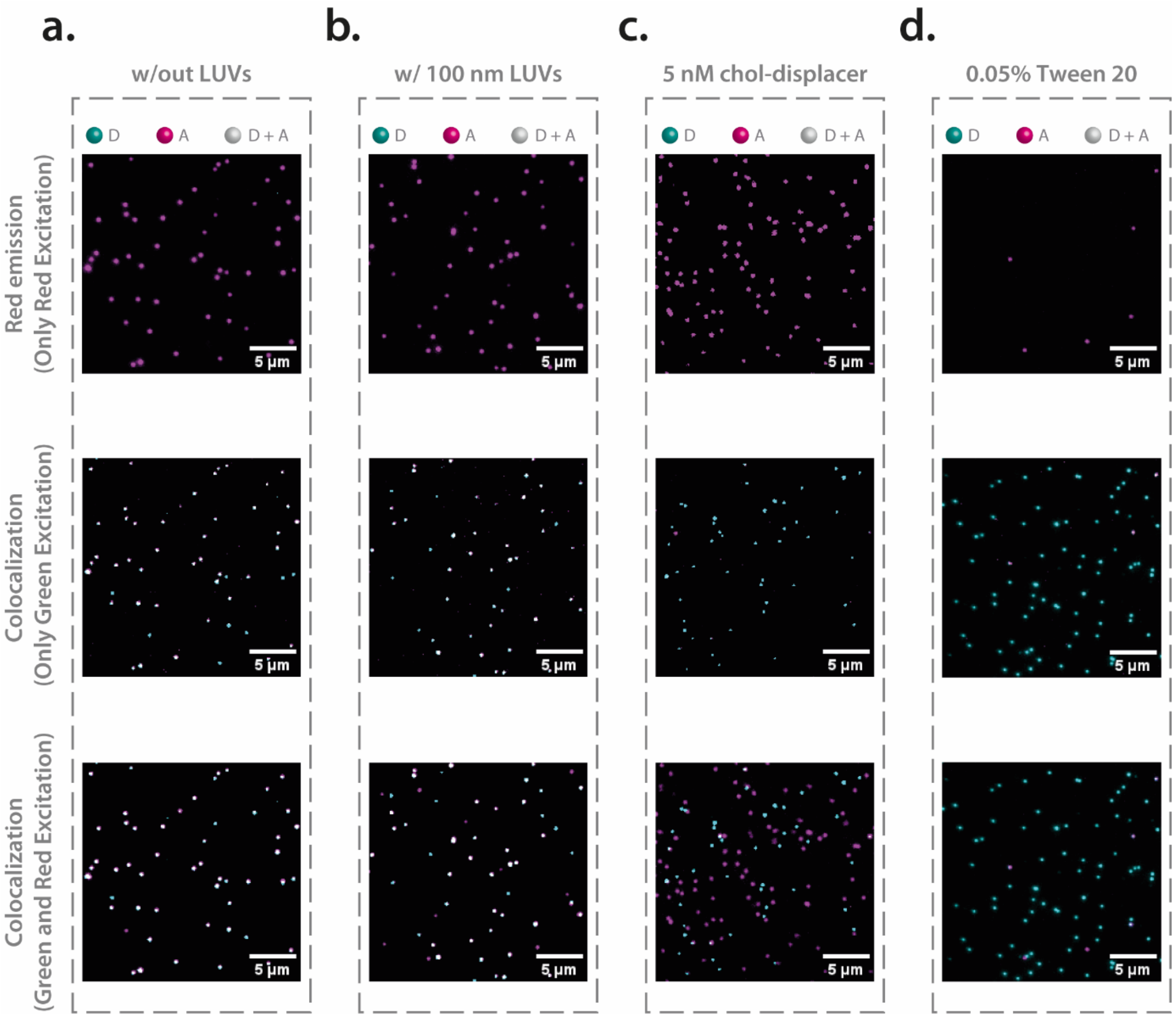

**Figure S4.** Validation of specific vesicle-probe interaction via the introduction of 0.05% Tween20. Each row presents distinct imaging modalities for different experimental stages: red emission after red excitation (top), superimposed TIRF images following green excitation (middle), and superimposed TIRF images after both green and red excitation (bottom). **(a)** Control system without lipid vesicles, serving as a baseline for fluorescence emission and colocalization. **(b)** System containing 100 nm lipid vesicles, demonstrating the initial state before displacement. **(c)** Post-incubation with a 5 nM cholesterol-labeled displacer strand, revealing the successful translocation of the ATTO647N-labeled probe to the vesicles. **(d)** Post-washing with a buffer containing 0.05% Tween20, showing a near-complete disappearance of red emission spots, which affirms the specificity of vesicle-probe interaction.

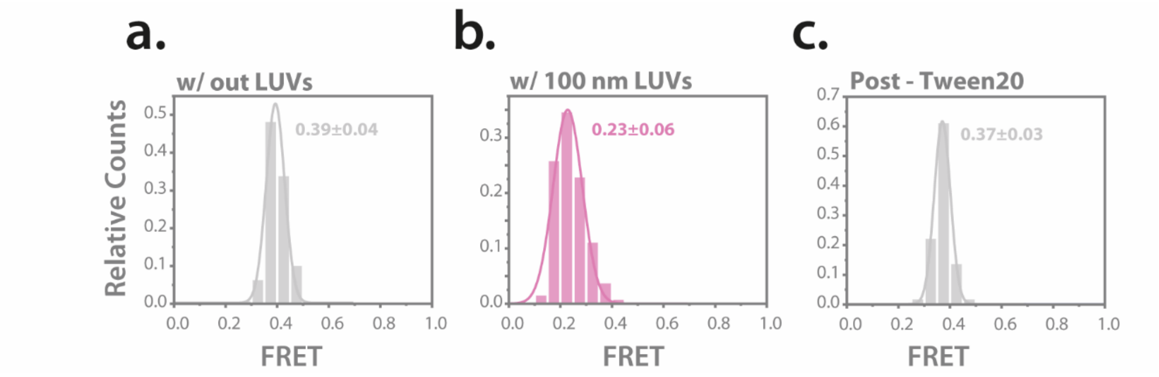

**Figure S5.** Examination of the impact of Tween20 on Sensor Behavior. The figure displays FRET histograms to evaluate the effect of 0.05% Tween20 on the system, specifically focusing on its ability to disrupt lipid vesicles without affecting sensor behavior. **(a)** FRET histogram of the system without lipid vesicles, serving as a control to establish baseline FRET values. **(b)** FRET histogram after the introduction of 100 nm lipid vesicles, demonstrating the change in FRET values upon vesicle binding. **(c)** FRET histogram following washing with a buffer containing 0.05% Tween20, but without prior incubation with the cholesterol-labeled displacer strand. The similarity between this histogram and the control (a) confirms that Tween20 exclusively disrupts the lipid vesicles without altering the FRET efficiency of the sensor. The error refers to the standard deviation (SD).

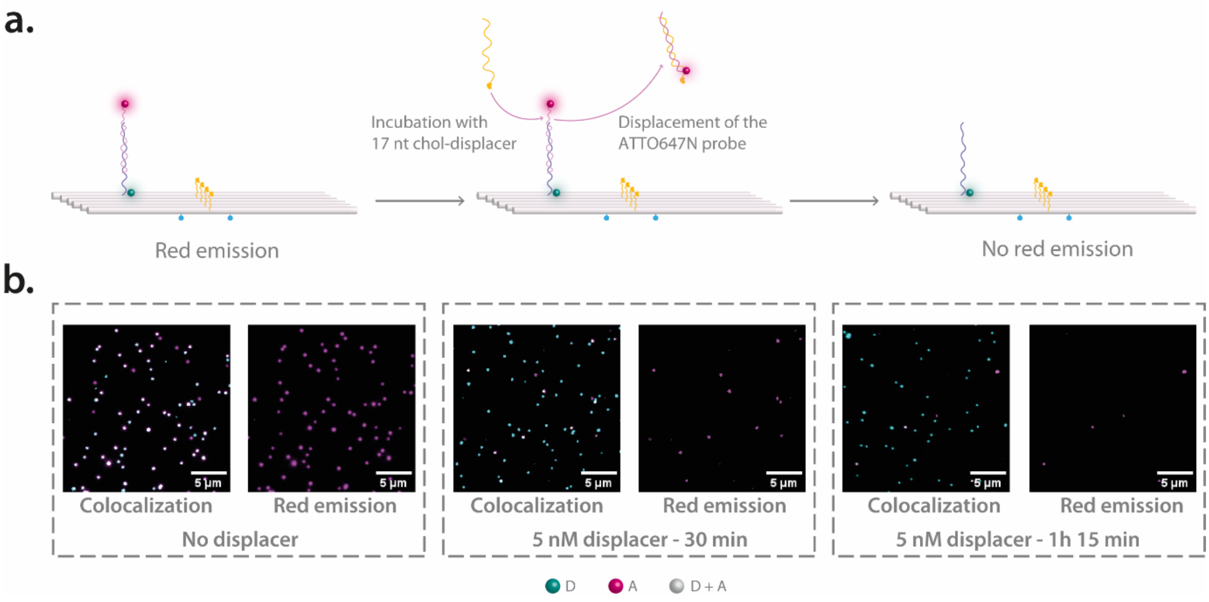

**Figure S6.** Experimental evidence illustrating the absence of non-specific probe adherence in a system devoid of lipid vesicles and the effectiveness of cholesterol-labeled displacer strand. **(a)** Schematic representation of the DNA origami-based sensor at various stages of the control experiment. The left panel shows the initial system with the ATTO647N-labeled sensing probe bound to the DNA origami, emitting red fluorescence. The middle panel illustrates the sensor after incubation with the 17 nt cholesterol-labeled displacer strand, depicting how the probe is displaced into solution due to the absence of lipid vesicles for anchoring. The right panel shows the sensor solely with the ATTO542 fluorophore, representing the disappearance of the red emission signal. **(b)** Superimposed TIRF images corroborating the above sequence of events. The left images show both colocalized and isolated red emission spots before the introduction of the cholesterol displacer strand. The middle images show reduced numbers of colocalized and red spots after 30 minutes of incubation with the displacer strand. The right images, taken after 1 hour and 15 minutes of incubation followed by washing, reveal a near-complete disappearance of red emission spots, confirming that the displacer efficiently translocates the sensing probe without unspecific sticking.

### 5. References

1. Douglas, S. M.; Marblestone, A. H.; Teerapittayanon, S.; Vazquez, A.; Church, G. M.; Shih, W. M., Rapid prototyping of 3D DNA-origami shapes with caDNAno. *Nucleic Acids Research* **2009,** *37* (15), 5001-5006.
2. Kapanidis, A. N.; Laurence, T. A.; Lee, N. K.; Margeat, E.; Kong, X.; Weiss, S., Alternating-Laser Excitation of Single Molecules. *Accounts of Chemical Research* **2005,** *38* (7), 523-533.
3. Lee, N. K.; Kapanidis, A. N.; Wang, Y.; Michalet, X.; Mukhopadhyay, J.; Ebright, R. H.; Weiss, S., Accurate FRET Measurements within Single Diffusing Biomolecules Using Alternating-Laser Excitation. *Biophysical Journal* **2005,** *88* (4), 2939-2953.
4. Margeat, E.; Kapanidis, A. N.; Tinnefeld, P.; Wang, Y.; Mukhopadhyay, J.; Ebright, R. H.; Weiss, S., Direct Observation of Abortive Initiation and Promoter Escape within Single Immobilized Transcription Complexes. *Biophysical Journal* **2006,** *90* (4), 1419-1431.

5. Vogelsang, J.; Kasper, R.; Steinhauer, C.; Person, B.; Heilemann, M.; Sauer, M.; Tinnefeld, P., A Reducing and Oxidizing System Minimizes Photobleaching and Blinking of Fluorescent Dyes. *Angewandte Chemie International Edition* **2008,** *47* (29), 5465-5469.
6. Cordes, T.; Vogelsang, J.; Tinnefeld, P., On the Mechanism of Trolox as Antiblinking and Antibleaching Reagent. *Journal of the American Chemical Society* **2009,** *131* (14), 5018-5019.
7. Preus, S.; Noer, S. L.; Hildebrandt, L. L.; Gudnason, D.; Birkedal, V., iSMS: single-molecule FRET microscopy software. *Nature Methods* **2015,** *12* (7), 593-594.
